# Red Panda: A novel method for detecting variants in single-cell RNA sequencing

**DOI:** 10.1101/2020.01.08.898874

**Authors:** Adam Cornish, Shrabasti Roychoudhury, Krishna Sarma, Suravi Pramanik, Kishor Bhakat, Andrew Dudley, Nitish K Mishra, Chittibabu Guda

## Abstract

Single-cell sequencing enables us to better understand genetic diseases, such as cancer or autoimmune disorders, which are often affected by changes in rare cells. Currently, no existing software is aimed at identifying single nucleotide variations or micro (1-50bp) insertions and deletions in single-cell RNA sequencing (scRNA-seq) data. Generating high-quality variant data is vital to the study of the aforementioned diseases, among others. In this study, we report the design and implementation of Red Panda, a novel method to accurately identify variants in scRNA-seq data. Variants were called on scRNA-seq data from human articular chondrocytes, mouse embryonic fibroblasts (MEFs), and simulated data stemming from the MEF alignments. Red Panda had the highest Positive Predictive Value at 45.0%, while other tools—FreeBayes, GATK HaplotypeCaller, GATK UnifiedGenotyper, Monovar, and Platypus—ranged from 5.8%-41.53%. From the simulated data, Red Panda had the highest sensitivity at 72.44%. We show that our method provides a novel and improved mechanism to identify variants in scRNA-seq as compared to currently-existing software.

**Availability:** Source code freely available under the MIT License at https://github.com/adambioi/red_panda, and is supported on Linux

## 1. Introduction

Single-cell sequencing (SCS) is a relatively new technique that saw its first use in 2011 (Navin et al., 2011) and has been used to investigate important biological problems: examining the heterogeneity of different cancers (Suzuki et al., 2015), determining copy number variation in enhanced detail (McConnell et al., 2013), and better characterizing circulating tumor cells using differential expression analysis (Ramsköld et al., 2012; Ni et al., 2013). Multiple recent studies using SCS have also shown that tumors are genetically diverse and produce subclones that contribute to the pathogenicity of the disease by conferring chemotherapy resistance and metastatic capabilities to the tumor (Gawad et al., 2014; Jan et al., 2012). This technology has also proven useful by aiding in characterizing somatic mutations in neurons (Lodato et al., 2015), identifying rare intestinal cell types (Grün et al., 2015), and discriminating cell types in healthy tissues (Jaitin et al., 2014; Zeisel et al., 2015).

One area that has not been widely explored is the detection of small variants in SCS. Single Nucleotide Variants (SNVs) and micro (1-50bp) insertions and deletions (indels) can have a large impact on human disease (Tennessen et al., 2012; Gill et al., 2014; Ku et al., 2013) and are typically identified using exome sequencing or whole-genome sequencing (WGS) (Lek et al., 2016). Monovar has been developed to identified variants in scDNA-seq (Zafara et al., 2016), but there exists no companion tool for scRNA-seq. An effort has been made to apply best practices for identifying variants in RNA-seq to scRNA-seq datasets (Lodato et al., 2015; Borel et al., 2015), but they do not take advantage of the unique nature of the data produced by the scRNA-seq platform.

This study introduces a novel method, Red Panda, that is designed specifically to identify variants in single-cell RNA sequencing (scRNA-seq) and tests how it compares to currently-available variant callers: FreeBayes (Garrison and Marth, 2012), GATK HaplotypeCaller (Poplin et al., 2017), GATK UnifiedGenotyper (DePristo et al., 2011), Platypus (Rimmer et al., 2014), and Monovar (Zafara et al., 2016). The first four tools were originally developed for calling variants using bulk DNA sequencing data but can also identify variants in bulk mRNA sequencing data. For our purposes, data from scRNA-seq, as opposed to scDNA-seq, is used as it largely avoids errors stemming from single-cell genomic sequencing—high allelic dropout, coverage nonuniformity leading to lack of coverage in exons, and False Positive (FP) amplification errors (Wang and Navin, 2015).

Red Panda employs the unique information found in scRNA-seq to increase accuracy as compared to software designed for bulk sequencing. We utilize the fact that transcripts represented by scRNA-seq reads necessarily only originate from the chromosomes present in a single cell. Where applicable, this fact is used to decide what is and is not a heterozygous variant. For example, if 20% of the transcripts of a gene originate from the maternal chromosome and 80% originate from the paternal, then all the heterozygous variants of that gene in the expressed transcript will be represented at a reference to alternate allele ratio of either 1:4 or 4:1. In other words, all the heterozygous variants in that transcript are expected to be part of a bimodal distribution, which can be exploited to improve the accuracy of variant calling using scRNA-seq data. Such unique information could not be obtained from bulk sequencing, where each variant is independently called. As part of the process of identifying variants, Red Panda creates three different classes: homozygous-looking, bimodally-distributed heterozygous, and non-bimodally-distributed heterozygous. We use simulated and experimental data to prove that this partitioning strategy, as well as treating bimodally-distributed variants differently, leads to an increase in sensitivity and Positive Predictive Value (PPV) compared to currently-available methods.

## 2. Methods

### 2.1. Data generation and quality control

For algorithm development, human articular chondrocytes were sequenced using the Smart-seq2 protocol for single cells (**Supplemental Figure 1**). These data satisfied five criteria needed for a test dataset: (i) bulk genomic sequencing data paired with scRNA-seq data generated from Smart-seq2 libraries, (ii) isogenic tissue, (iii) high quality sequencing data, (iv) the sequencing data must be from an organism with a well-annotated genome, (v) the sequencing data must come from normal cells. The first criterion was especially important because the bulk sequencing data was used to corroborate the findings from the scRNA-seq data. For this dataset, 30 live cells successfully captured from a 96 chamber C1 Fluidigm IFC were sequenced and eight were removed due to: low read count, too many reads originating outside exons (percentage of reads outside exons is one standard deviation above the median percentage of reads outside exons for all samples), and/or transcription profiles not correlating with the other cells sequenced (Pearsons Correlation: p>0.05; **Supplemental Figures 2 & 3 and Supplemental Table 1**).

**Table 1.** PPV and FDR for each tool. The average PPV and FDR with standard deviations for each tool using the exome as reference is listed.

| Algorithm | Average PPV (%) | Average FDR (%) |
| --- | --- | --- |
| FreeBayes | 8.69% ± 0.35% | 91.31% ± 0.35% |
| GATK HaplotypeCaller | 31.67% ± 2.08% | 68.33% ± 2.08% |
| GATK UnifiedGenotyper | 5.84% ± 0.45% | 94.16% ± 0.45% |
| Monovar | 41.53% ± 0.29% | 58.47% ± 0.29% |
| Platypus | 6.95% ± 0.49% | 93.05% ± 0.49% |
| Red Panda | 44.96% ± 3.15% | 55.04% ± 3.15% |

**Figure 1.**
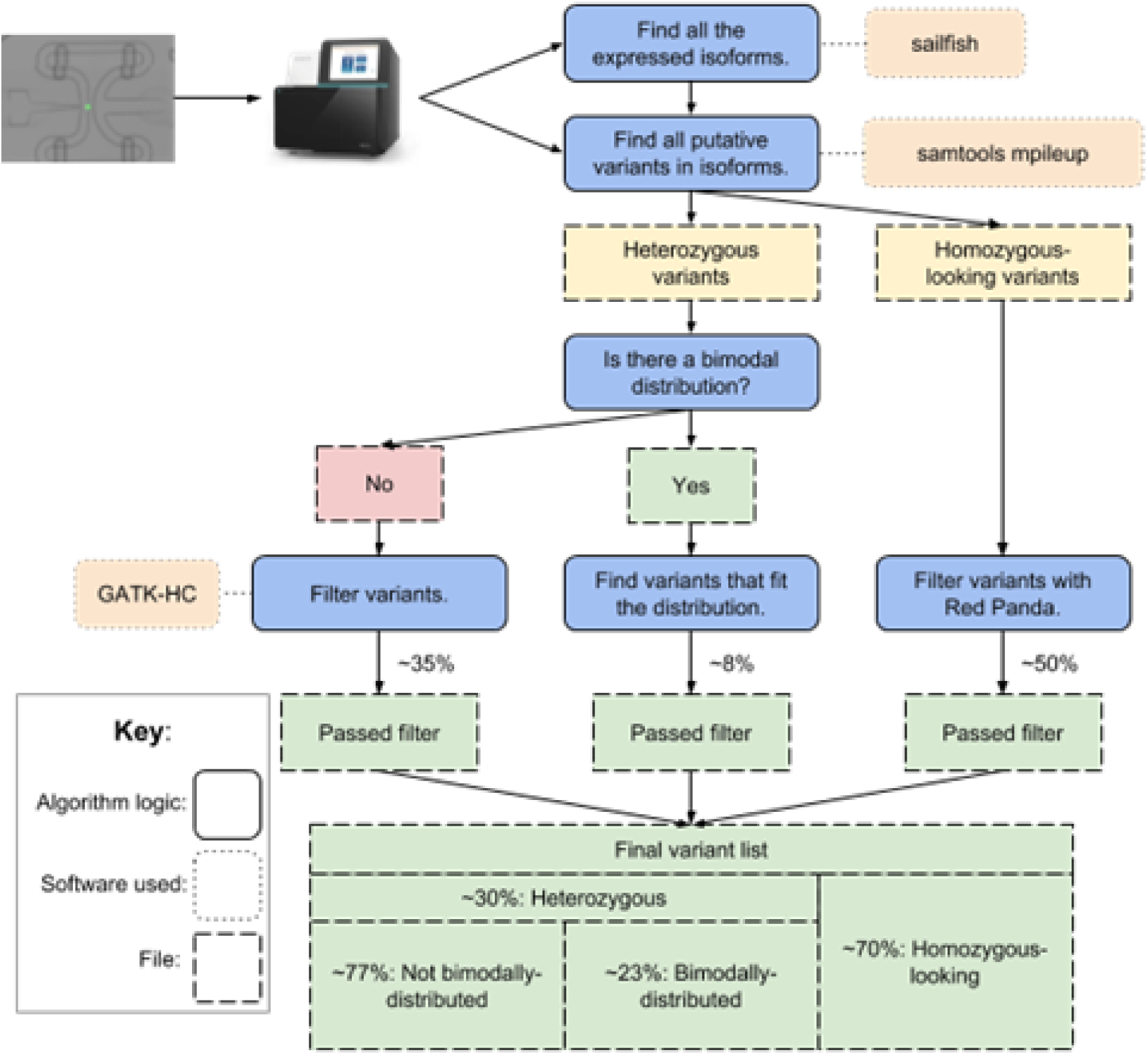
A simple schematic of the logic used in Red Panda. For every cell, every expressed isoform is identified with sailfish. All putative variants are then identified in each isoform and split into a homozygous-looking VCF file and a heterozygous VCF file. Then the former is filtered by Red Panda using quality cutoffs while the **latter is filtered using Red Panda if the variants are bimodally-distributed or GATK-HC if they are** not. These three sets of variants are then combined into a single VCF file.

**Figure 2.**
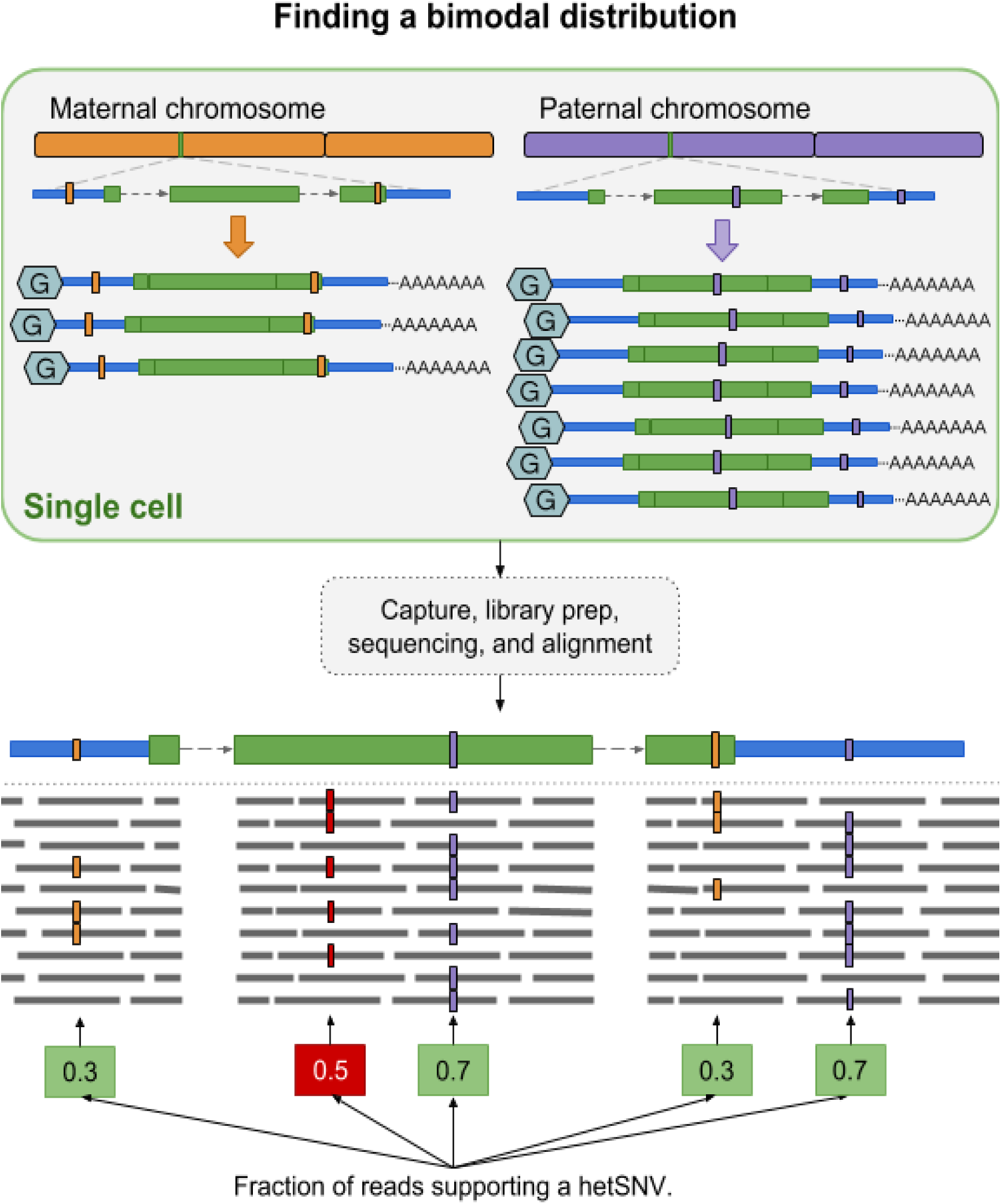
Finding a bimodal distribution. Any variants (green box) that fit into the expected distribution of reads stay. Any that do not are removed: here the variant existing at a fraction of 0.5 (red box) would be removed.

Additionally, 56 mouse embryonic fibroblasts (MEFs) were sequenced using the Smart-seq2 protocol and are paired with Sanger sequencing for validation. Simulated data were generated from the MEF alignment files for each sample (**Supplemental Figure 4**). Of these, one cell was removed due to its low read counts, too many reads originating outside exons, and transcription profiles not correlating with the other cells sequenced (**Supplemental Figures 5 & 6**).

### 2.2. Exome sequencing

We performed cell prep and DNA extraction on human articular chondrocytes harvested the same day from the same batch as the single-cell capture. Genomic DNA was extracted using the QIAGEN DNA extraction kit per manufacturer’s instructions. Due to the low amount of DNA captured (80ng), 12 PCR amplification cycles were performed prior to library preparation to obtain enough DNA. The Agilent SureSelect Clinical Research Exome V2 kit was used to capture coding regions and generate a library. The exome library was sequenced on two lanes of the NextSeq500 using 75 base pair paired-end sequencing.

### 2.3. Exome variant calling

The bcbio-nextgen v. 1.0.3 pipeline was used for variant calling to align reads and identify variants in the exome. Reads were aligned to the human genome v. 38 (hg38) using BWA-MEM v. 0.7.15. FreeBayes (v. 1.1.0), GATK HaplotypeCaller (v. 3.7.0), and Platypus (v. 0.8.1) were used to identify SNVs and indels. Only those variants identified by at least two out of the three algorithms were kept. MultiQC v. 1.0.dev0 was run to aggregate Quality Control (QC) statistics from bcbio-nextgen, samtools v. 1.4, bcftools v. 1.4, and FastQC v. 0.11.5.

### 2.4. Single-cell RNA Sequencing

#### 2.4.1 Human articular chondrocytes

Articular chondrocytes were harvested from a Caucasian female patient undergoing total knee replacement, who provided informed consent under IRB #691-13-EP prior to the study. Cells were extracted from shavings of articular cartilage, all of which was consumed in the generation of the scRNA-seq and exome libraries. This was done through sequential digestion in .2% Pronase (Roche) for 2 hours followed by overnight digestion in .2% collagenase (Gibco), all while shaking at 37°C. Cell suspensions were passed through 70μM cell strainers (BD Falcon) and centrifuged at 500xG for 10 minutes to recover chondrocytes. The cells were subsequently embedded in three-dimensional alginate bead cultures at a final concentration of about 75 million cells per mL. The cultures were maintained at 37°C in a 5% CO2 atmosphere in Dulbecco’s modified Eagle medium (DMEM)/F12 (1:1) supplemented with 1% penicillin-streptomycin-glutamine (Invitrogen, 10378-016), Amphotericin B (Gibco, 15290026), insulin-transferrin-sodium selenite (Sigma, I2771), 50μg/mL Vitamin C, 10ng/mL FGF2, and 10ng/mL TGF-bb3 (PeproTech®, 100-36E) for 14 days. The day before single-cell capture, cells were lysed using Trizol® reagent (Life Technologies) according to the manufacturer’s protocol. These cells were split into two groups for DNA and RNA extraction. Cells were loaded onto a 10-17 μm Fluidigm C1 Single-Cell Auto Prep IFC, and the cell-loading script was performed using the manufacturer’s instructions. Each of the 96 capture sites was inspected under a confocal microscope to remove sites containing dead cells (as identified by the LIVE/DEAD Cell Viability Assay) and to remove capture sites containing more than one cell. Cells that were not identified as either alive or dead by the LIVE/DEAD assay were retained for RNA sequencing. Following capture, reverse transcription and cDNA amplification were performed in the C1 system using the Clontech SMARTer Ultra Low Input RNA Kit for Sequencing v3 per the manufacturer’s instructions. Amplification was performed using the Nextera XT DNA Sample Preparation Kit, and the Nextera XT DNA Sample Preparation Index Kit (Illumina) was used for indexing. After quantification using an Agilent Bioanalyzer, sequencing was performed on two lanes of the NextSeq500 using 150 base pair paired-end sequencing.

#### 2.4.2 Mouse embryonic fibroblasts

Mouse embryonic fibroblasts (MEFs) were harvested from embryos at E13.5 and extracted using previously standardized methods (Xu, 2005). After isolation, cells were cultured in DMEM containing 10% FBS and 1% of each penicillin and streptomycin at 37°C in a 5% CO2 atmosphere for 2 days. On the day of single-cell capture, cells were trypsinized (0.05% Trypsin-EDTA solution), counted, and resuspended in media at 105 cells/mL concentration. Sequencing was performed as described above.

### 2.5. Single-cell RNA variant calling

The bcbio-nextgen v. 1.0.3 pipeline for RNA-seq was used to align reads and perform transcript quantification for each cell. Reads were aligned using hisat2 v. 2.1.0 to be used in the downstream analysis for FreeBayes, GATK HaplotypeCaller, GATK UnifiedGenotyper, Monovar, and Red Panda. However, for Platypus, BWA-MEM v. 0.7.15 was used to align reads due to Platypus’s inability to process reads split across long distances. The genome hg38 was used for the human articular chondrocytes, and mm10 was used for the MEFs. All four bulk variant callers were run using default parameters with a few exceptions. For FreeBayes, min-alternate-fraction was set to 0.1 and no-partial-observations was enabled, and GATK HaplotypeCaller and GATK UnifiedGenotyper set standard minimum confidence threshold for calling to 4.0. Sailfish v. 0.10.1 was used to generate expression values. MultiQC v. 1.0.dev0 was run to aggregate QC statistics from bcbio-nextgen, samtools v. 1.4, QualiMap v. 2.2.2a53, and FastQCv. 0.11.5.

### 2.6. Sanger sequencing

Primer3Plus (**Supplemental Table 2**) was used to create target regions for Sanger sequencing. For the first round of sequencing, the PCR reaction was performed using GoTaq Hot Start Polymerase following the manufacturer’s protocol with an annealing temperature (Tm) of 50°C. After amplification, PCR products were run on 1.5% agarose gel and visualized in Kodak gel doc, and specific DNA bands were recovered using QIAquick Gel Extraction Kit. For the second round of sequencing of the Red Panda-specific variants, a 55°C Tm was used, followed by running the PCR products on 2% agarose gel. Purified DNA products paired were submitted to Genewiz for Sanger sequencing.

**Table 2.** Average variant count and standard deviation for each tool. For this analysis, the total number of variants identified by each tool is reported.

|  | FreeBayes | GATK-HC | GATK-UG | Monovar | Platypus | Red Panda |
| --- | --- | --- | --- | --- | --- | --- |
| Average | 865.8 | 611.1 | 574.7 | 3515.44 | 315.4 | 1071.8 |
| Stdev | 235.3 | 195.1 | 170.5 | 834.04 | 107.1 | 372.6 |

### 2.7. Statistic calculations

For our calculations, (i) a True Positive (TP) is a position on the genome that is correctly identified as differing from the reference genome, (ii) a True Negative (TN) is a position on the genome that is correctly identified as not differing from the reference, (iii) a False Positive (FP) is a position on the genome that is incorrectly identified as differing from the reference, (iv) a False Negative (FN) is a position on the genome that is incorrectly identified as not differing from the reference genome.

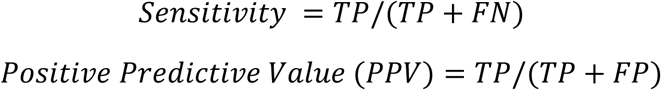

### 2.8. Software requirements and distribution

Red Panda, written almost entirely in Perl, relies on samtools mpileup and GATK HaplotypeCaller to function. The tool mpileup is required to generate a list of every variant in a sample. GATK HaplotypeCaller is necessary to call heterozygous variants that do not fit a bimodal distribution. Bedtools, vcf-sort found in the vcftools package, and Picard Tools are all necessary to manipulate the different types of files used during the variant calling process. As these tools are all supported by different institutions under different licenses, Red Panda does not come prepackaged with them. Red Panda is released under the MIT license and can be found on GitHub at https://github.com/adambioi/red_panda.

## 3. Algorithm

### 3.1 Red Panda

Red Panda takes two files as input: a tab-delimited file generated by sailfish (Patro *et al*., 2014) containing a list of all isoforms and their expression levels in a cell and also a Variant Call Format (VCF) file generated by samtools mpileup (Li, 2011) containing a pileup of all locations in the cell’s genome that differ from the reference. The second file is the list of all putative variants from which Red Panda will create three classes: homozygous-looking, bimodally-distributed heterozygous, and non-bimodally-distributed heterozygous variants.

The distinction between heterozygous variants and homozygous-looking variants is necessary, because variants will either have a fraction of reads that support an unambiguously heterozygous variant, or they will have a fraction of reads that, in a single cell, appears to be a homozygous variant, but could potentially be heterozygous. This is due to the stochastic nature of RNA transcription leading to allele-specific expression (Yan, 2002; Gregg *et al*., 2010; Marinov *et al*., 2014). This monoallelic expression can lead to a heterozygous variant looking like a homozygous variant (Deng *et al*., 2014; Gimelbrant *et al*., 2007). Due to this ambiguity, variants that have full read coverage supporting an alternate allele are hereafter termed “homozygous-looking” rather than “homozygous”. The workflow for this methodology can be found in **Figure 1**.

Red Panda capitalizes on the fact that data come from a single cell, so transcripts represented by the scRNA-seq reads necessarily come from the two chromosomes present, and we factor that into our decision-making process when establishing what is and is not a variant. In a diploid cell, one would expect transcripts to originate from two chromosomes, and thus, any heterozygous variant present in a given transcript will be represented in the sequencing data in a fraction consistent with the fraction of transcripts coming from a specific chromosome. **Figure 2** shows that if 30% of the transcripts for a gene in a cell originate from the maternal chromosome and 70% from the paternal chromosome, then reads in the scRNA-seq data will represent every heterozygous variant present in that transcript at either a 7:3 ratio (reference: alternate allele) or a 3:7 ratio. This type of variant is considered to be bimodally-distributed heterozygous and any variant on the same transcript that’s falling outside of this distribution with a tolerance of 5% is likely to be a False Positive. Using this concept, Red Panda can accurately remove False Positive heterozygous variant calls—often artifacts from the library preparation, sequencing, or alignment—as well as identify variants supported by even a low fraction of reads that the current tools would not be able to capture (**Supplemental Figure 7**).

The VCF file generated by samtools mpileup is split into two lists containing variants that are heterozygous or homozygous-looking. Heterozygous variants are filtered into files containing bimodally-distributed and non-bimodally-distributed variants, the latter of which is filtered by GATK HaplotypeCaller as there is no unique information that Red Panda can capitalize on and it has been proven to be among the most accurate variant callers available (Cornish and Guda, 2015). The final list of variants is bimodally-distributed heterozygous, heterozygous not fitting a bimodal distribution but supported by GATK HaplotypeCaller, and homozygous-looking. This method of partitioning variants is also used for indels.

Red Panda runs on a single core but can easily be parallelized by being run on a cluster. Each cell, each with ∼5.4 million reads, takes, on average, two hours to complete the analysis.

### 3.2 Simulation

Roughly 1,000 simulated variants were programmatically inserted into the alignments generated from the MEFs resulting in a unique set of simulated variants for each cell. Of these, 650 were homozygous, and ∼350 were heterozygous, a subset of which ∼70 were bimodally-distributed. These numbers were used because they are close to the proportions seen in the variants corroborated by the exome sequencing in the articular chondrocyte data, and while these proportions do not match those expected based on bulk sequencing experiments (Dewey *et al*., 2016; 1000 Genomes Project Consortium *et al*., 2010; Choi *et al*., 2009), they do match what is expected from scRNA-seq data (Borel *et al*., 2015). To instill a level of uniformity, variants were only inserted where there was at least 20x read coverage (**Supplemental Figure 8**). The positions on the genome where the 650 homozygous and 280 non-bimodally-distributed heterozygous variants were inserted were not restricted except that they must originate from the locations with at least 20x read coverage.

The bimodally-distributed heterozygous variants had additional parameters determining their placement. They were required to have a minimum of two variants placed in an expressed isoform. From the MEF sequence data, an average of ∼3 (a range of 2–5) variants per isoform were observed resulting in 23 randomly chosen genes being used for this class of variant. For each isoform, 2–5 variants were randomly inserted into the gene but only if more than 250bp of viable (read depth >= 20x) locations existed.

## 4. Results

### 4.1 Comparison of different tools using human articular chondrocytes

Alignment files generated for each of the 22 chondrocyte scRNA-seq samples were used as input for FreeBayes, GATK HaplotypeCaller, GATK UnifiedGenotyper, Monovar, Platypus, and Red Panda. The variant calls generated by each tool were cross-referenced with the variants found in the exome to determine their veracity. To avoid False Negatives, comparisons were restricted to locations supported by alignments in both the exome and the cell being compared. To evaluate the ability of each method, the number of variants found in concordance with the exome as well as the PPV for each tool were calculated.

**Figure 3a** shows that, on average, Red Panda identifies 913 variants per cell that are in accordance with the exome whereas FreeBayes identifies 65, GATK HaplotypeCaller 705, GATK UnifiedGenotyper 222, Monovar 861, and Platypus 386. There is a consistent overlap between the tools, even for FreeBayes and GATK UnifiedGenotyper which typically did not identify as many variants as the other tools (**Supplemental Figure 9**). While Red Panda shares significant overlap with the other tools, it also identifies a large number of unique variants by itself.

**Figure 3.**
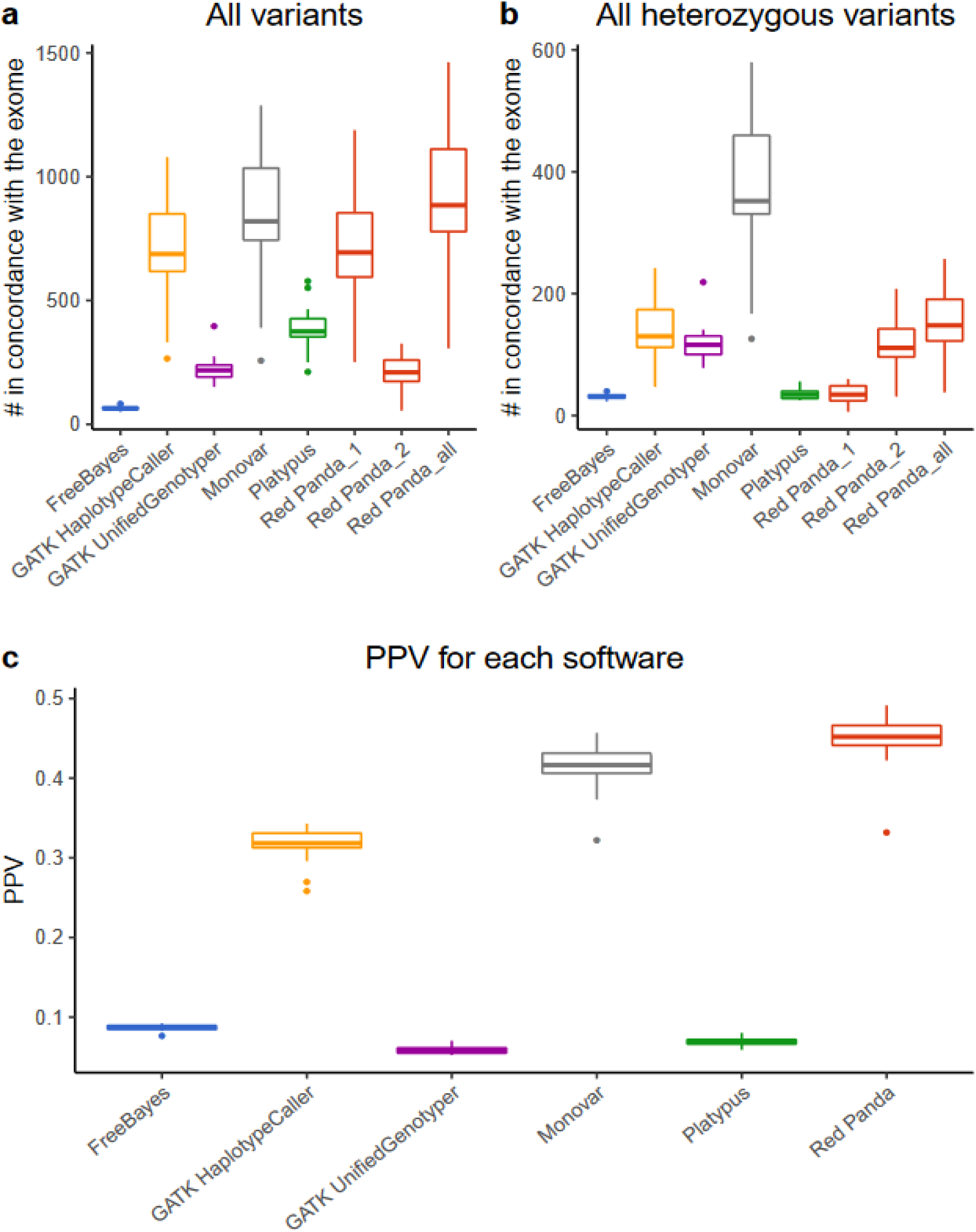
Variants in concordance with the exome and PPV for each tool. **(a)** The total number of variants in concordance with the exome for every cell as identified by each tool. Red Panda is characterized by three box plots: 1, 2, and all. Red Panda_1 contains variants exclusive to Red Panda logic: homozygous-looking variants and bimodally-distributed heterozygous variants. Red Panda_2 contains non-bimodally-distributed heterozygous variants that are called by GATK-HaplotypeCaller. Red Panda_all is a superset of the two. **(b)** The total number of heterozygous variants in concordance with the exome for every cell as identified by each tool. Red Panda is characterized by three box plots: 1, 2, and all. Red Panda_1 contains bimodally-distributed heterozygous variants. Red Panda_2 contains non-bimodally-distributed heterozygous variants. Red Panda_all is a superset of the two. **(c)** The average PPV calculated for each tool.

To assess the effectiveness of these variant callers with regards to heterozygous variant identification, the same analysis was performed using just the heterozygous SNVs and indels in each sample. Each tool has its own annotation dictating if a variant is heterozygous, and these variants were cross-referenced with the exome sequencing data to confirm the variants were, in fact, heterozygous. **Figure 3b** shows the total number of heterozygous SNVs and indels in concordance with the exome for each tool and each cell. On average 154 variants in agreement with the exome were identified by Red Panda, 31 by FreeBayes, 136 by GATK HaplotypeCaller, 118 by GATK UnifiedGenotyper, 368 by Monovar, and 36 by Platypus. PPV and False Discovery Rate (FDR) were calculated (**Table 1** and **Figure 3c**), and show that Red Panda has the highest average PPV (44.96%) of any of the tools.

### 4.2 Comparison of different tools using MEFs

The six software packages were compared by assessing the variant overlap between cells. As these cells were isogenic, each cell should have shared a large portion of its variants with the other cells sequenced. This was evaluated with a high overlap identified by a variant caller as an indicator that that software performed well.

**Table 2** shows the average number of variants identified with Monovar identifying the highest number of variants of all the tools. Extrapolating from the PPV results from the articular chondrocyte data, this means that Monovar also identified the highest number of True Positives and fewest False Positives.

Overlap of variants between cells was measured by looking at the all-to-all comparison of 55 cells, resulting in 1,540 unique possible comparisons. Three groups of variants are assessed in these comparisons: all variants shared, homozygous-looking variants shared, and heterozygous variants shared (**Supplemental Figures 10 & 11**).

**Figure 4** shows the distribution of the fraction and a total number of variants overlapping in the pairwise comparisons for all three classes of variants. Monovar identifies both the highest fraction of variants shared in pairwise comparisons as well as total variants shared. For Red Panda, the highest fraction and total count of variants shared in pairwise comparisons come from the homozygous-looking class wherein more than 75% of the comparisons achieve a higher fraction of overlap than every other tool. Interestingly, it is rare for any tool to have more than 100 heterozygous variants shared between cells.

**Figure 4.**
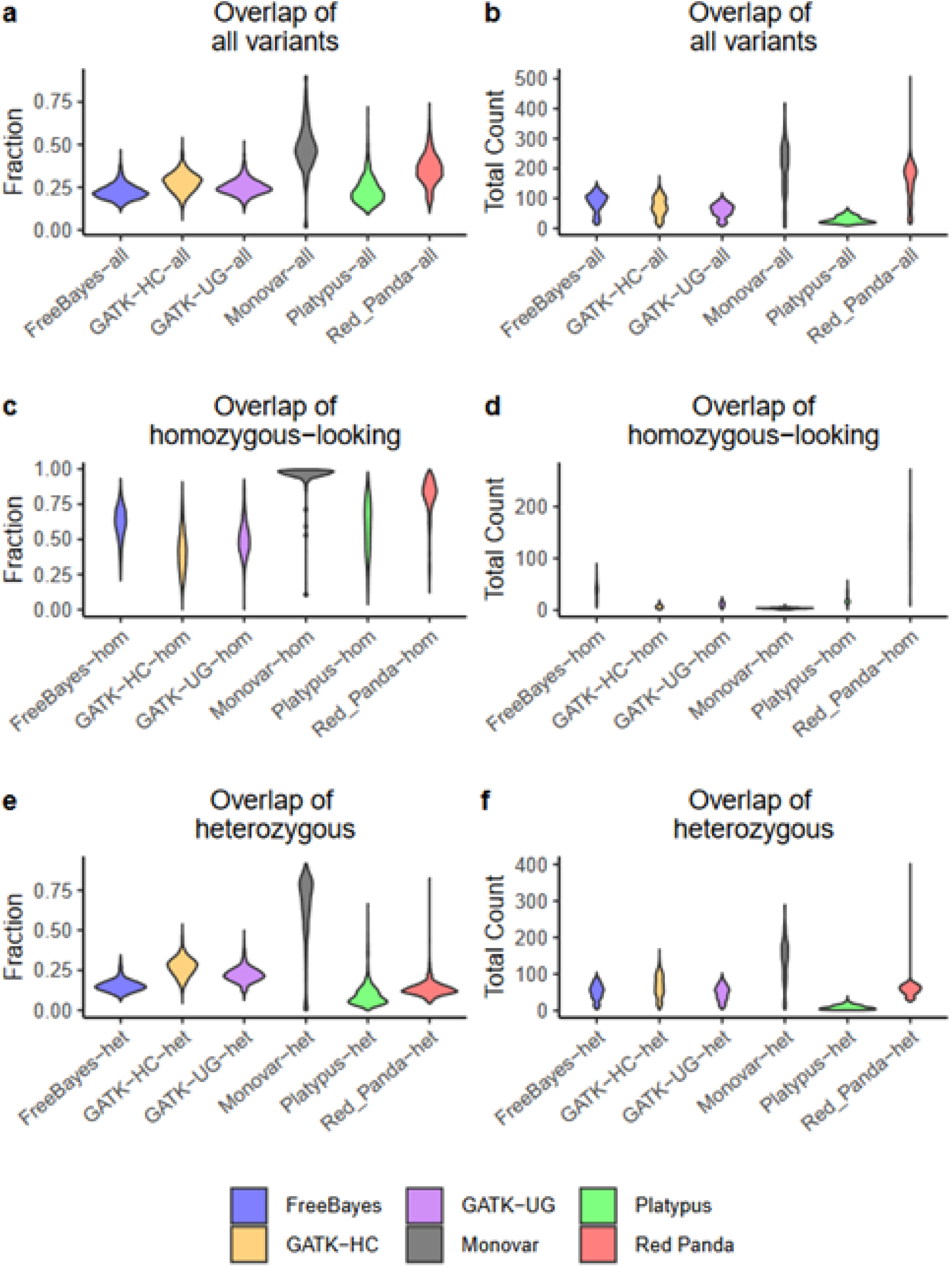
Violin plots for variants shared between cells. Violin plots show the fraction (left) and quantitative (right) overlap for (a, b) all variants, (c, d) homozygous-looking variants, and (e, f) heterozygous variants shared in every pairwise cell comparison.

### 4.3 Validation with simulated data

Sensitivity was calculated for each tool across every cell. **Figure 5** and **Supplemental Figure 12** show that for homozygous variants and bimodally-distributed heterozygous variants, Red Panda consistently outperforms the other four tools, resulting in a higher overall sensitivity. For heterozygous variants taken as a whole, Monovar performs the best of the tools. It is unsurprising then that, compared to Monovar, Red Panda does not perform as well in this category because it uses GATK HaplotypeCaller (shown to accurately only a identify few heterozygous variants in this simulation) to validate heterozygous variants that do not follow a bimodal distribution. In this instance, GATK HaplotypeCaller and GATK UnifiedGenotyper perform poorly because they both utilize a feature that considers all samples simultaneously. This results in inferior performance on a group of samples where each sample may have a large number of mutations unique to that sample, and for this simulation, every cell has a ∼1,000 variants unique to it. Red Panda does not suffer as much from this limitation as it explicitly directs GATK-HC to call variants at specific locations one at a time rather than jointly. However, this can result in lowered sensitivity for Red Panda as compared to Monovar on samples that are genetically similar where the latter identifies the highest fraction (**Figure 4e**) and highest total number (**Figure 4f**) of heterozygous variants shared in pairwise comparisons.

**Figure 5.**
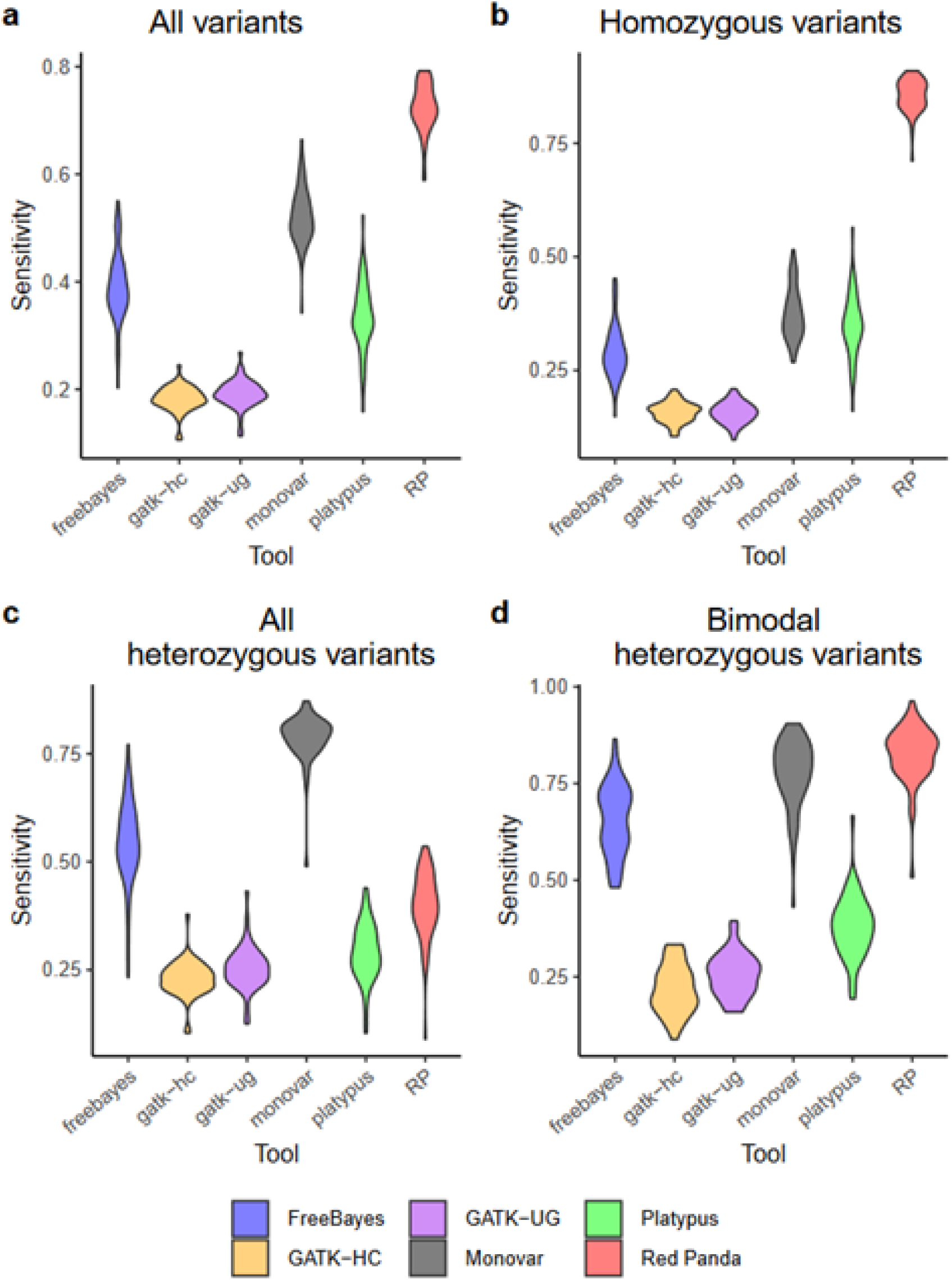
Sensitivity for identifying simulated variants for each tool. The violin plots of the sensitivity, calculated for each cell using each class of simulated variants are shown: **(a)** all variants, **(b)** homozygous variants, **(c)** all heterozygous variants, and **(d)** bimodally-distributed variants.

### 4.4 Validation with Sanger sequencing

Sanger validation was performed on two sets of random variants found in the MEF sequencing: one set of 20 random variants identified by all variant callers, and one set of 20 random variants identified exclusively by Red Panda. The first group is meant to assess the accuracy of all the tools taken as a whole, and the second is to address whether the Red Panda-specific variants are reliable. One requirement of the variants being validated is that they were identified in at least two cells. Ideally, variants present in more than 50% of the cells would be chosen, but as **Table 3** shows, there were not enough variants that are present in even >22 out of 55 of the cells to perform validation in this way. Enough valid sequences were generated for 33 of the 40 targets by Sanger sequencing to validate the presence of their corresponding variant (**Supplemental Tables 7–9**). Of these 33, only three variants, all of which were exclusively identified by Red Panda, were validated by Sanger sequencing. In all three instances, the variants were found in nine or more cells.

**Table 3.** Breakdown by tool of variants present in more than one cell. The number of cells in which a variant was found was broken down into five groups: presence in at least 2, 5, 10, 23, or 42 of cells. Additionally, the variants identified by all tools were checked for their presence in the five groups listed above. The variants submitted for Sanger sequencing were drawn from the two groups labeled with a cross (†) and an asterisk (*).

| Present in: | FreeBayes | GATK-HC | GATK-UG | Platypus | Red Panda | Intersection of all tools |
| --- | --- | --- | --- | --- | --- | --- |
| ≥ 2/55 cells | 2922 | 2463 | 1991 | 2947 | 3159† | 96* |
| ≥ 5/55 cells | 970 | 894 | 693 | 324 | 1051 | 18 |
| ≥ 10/55 cells | 416 | 398 | 309 | 161 | 565 | 0 |
| ≥ 23/55 of cells | 129 | 122 | 84 | 66 | 257 | 0 |
| ≥ 42/55 of cells | 38 | 24 | 27 | 22 | 98 | 0 |

## 5. Discussion

Identification of SNVs and indels is vital in addressing biological problems with a genetic component. While variant calling methods exist for samples collected from bulk sequencing, it is also important to have methods designed for samples collected from SCS. As shown with the experimental and simulated data, Red Panda makes it possible to perform variant detection in scRNA-seq with higher accuracy as compared to currently available software. Red Panda gains an advantage against other tools by intentionally separating variants into three separate classes and processing them differently: homozygous-looking, bimodally-distributed heterozygous, and non-bimodally-distributed heterozygous.

Using the exome variant data from bulk sequencing as a reference, Red Panda outperforms the other software. It provides both the highest PPV (45% -**Table 1**) of any of the tools as well as the highest number of variants in concordance with the exome (913 on average -**Figure 3a**). However, PPV is still low compared to using bulk sequencing data (Cornish and Guda, 2015; Sandmann et al., 2017).

In the MEF sequencing analysis, Red Panda performs better than the four bulk variant callers by identifying the highest number of variants per cell, but does fall short of the total identified by Monovar (**Table 2**). This could be explained by the fact that Red Panda is designed to work solely on data from individual cells, while Monovar gains an advantage by using variant data from all cells in the sample to calculate a posterior probability for making more confident variant calls. Surprisingly, both Monovar (3,515) and FreeBayes (865) identified a much higher number of variants (**Table 2**) as compared to the results from the human articular chondrocyte data (**Figure 3a**). The FreeBayes results are especially unexpected where it had the fewest number of variants shared between the scRNA-seq results and the exome (**Figure 3a**). One explanation may be that while FreeBayes identifies a high number of variants, the majority of those are False Positives. This idea is supported by the PPV for FreeBayes from the articular chondrocyte data (**Figure 3c**).

The pairwise cellular comparisons (**Figure 4**) assessed whether each variant caller performed well based on the consistency of their calls or if they performed poorly, randomly identifying variants in each cell, because, presumably, the same variants should exist in all 55 datasets. Red Panda performs extremely well for homozygous-looking variants, but is average for heterozygous variants when compared to the other tools, especially Monovar. This is due to the fact that, while Red Panda in principle confers an algorithmic advantage to identifying heterozygous variants, the monoallelic nature of gene expression and uneven sequencing coverage depth may preclude the tool from realizing its full potential. The majority of the heterozygous variants identified are actually evaluated by GATK HaplotypeCaller, since most are unsupported by a bimodal distribution (**Figure 3b**).

The results from the raw number and the fraction of variants overlapping in these pairwise comparisons show that Red Panda has higher distributions in both categories. It is possible to have a large number of variants shared, but also have a smaller fraction of variants shared between cells as is the case with FreeBayes. This indicates that there are more potential False Positives in the data generated by FreeBayes which fits with what was seen in the articular chondrocyte data (**Figure 3c**). The high fraction of homozygous-looking variants shared (**Figure 4c**) makes sense as it is less likely that allelic dropout will occur in this class as a result of allele-specific expression making for a more stable population of variants in the scRNA-seq data. Additionally, it is likely that it is rare for any tool to have more than 100 heterozygous variants shared between two cells because of the stochastic nature of allele-specific expression.

It is clearly possible to detect significant variation in transcripts at the single-cell level, but validating those variants at the DNA level is a big challenge due to the inability to isolate sufficient quantities of DNA from the same single cell to perform Sanger sequencing. Hence, we relied on Sanger sequencing performed on a bulk of DNA originating from the population of cells from which we harvested the single cells, which validated only three out of 33 variants tested. It is possible that the variants that failed validation are: False Positives, private mutations to a very small subset of cells that couldn’t be captured in the bulk DNA used from the population of cells, or they could be errors introduced in the scRNA-seq library preparation process. Due to these limitations, we relied on the simulated data for MEF cells and paired genomic sequencing as done with the human articular chondrocytes.

There are a number of methods available to imitate read counts and expression profiles (Zappia *et al*., 2017; Risso *et al*., 2018; Severson *et al*., 2018) in scRNA-seq; however, there currently exist no tools to generate scRNA-seq reads *in silico*, making the type of simulation carried out in this study necessary. We created a more realistic environment because the artifacts and flaws inherent to scRNA-seq are considered and maintained by our code. One downside however, is that the dataset used disallows calculating any accuracy statistics requiring False Positive numbers such as specificity and PPV, as the scRNA-seq also contains real variation inherent to the MEFs that will be picked up by each variant caller.

Based on the results from the simulated data (**Figure 5**), we found that Red Panda proves its advantage in identifying bimodally-distributed variants as well as homozygous variants, a class of variant that saw other tools struggle. When assessing total heterozygous variants, Monovar is superior to the other tools. This is somewhat surprising as Monovar gains a large part of its advantage in pooling cells together to identify variants, a strategy that should afford it no advantage given our method of simulating variants (each cell was assigned 1000 variants unique to that cell). Another unexpected result was how well FreeBayes performed given what was seen in the results from the human articular chondrocyte experiment where FreeBayes identified very few variants in concordance with the list obtained from exome sequencing, but a clue as to why this is might be found in the results from GATK-HC and GATK-UG. Both of these perform similarly to each other in the simulation with the latter consistently performing slightly better than the former. When variants were called in the exome to generate a reference list against which these variants from the scRNA-seq data could be compared, variants were only retained if they were identified by at least two of the following three tools: FreeBayes, GATK-HC, and Platypus. However, if the variants in this reference list were consistently only supported by the latter two, then it follows that variants identified by FreeBayes in the scRNA-seq experiment would be filtered out and make it appear as though FreeBayes identified a low number of True Positives. This is further supported by the fact that Platypus identifies many more variants in concordance with the exome than FreeBayes. The simulated data indicate that FreeBayes has good sensitivity, but identifies a large set of variants different from both GATK-HC and Platypus. Given the above, in order to improve the accuracy of Red Panda, Monovar or a combination of Monovar and GATK-HC could be used when evaluating non-bimodally-distributed heterozygous variants.

It is important to note that the advantages conferred by Red Panda are currently limited to scRNA-seq generated by library-preparation methods that generate full-length transcripts from cDNA such as Smart-seq2 or Holo-seq (Xiao et al., 2018), although the latter has not been tested in this study. As methods such as G&T-seq (Macaulay *et al*., 2015) mature—G&T-seq produces genomic and transcriptomic sequencing from the same cell—Red Panda can be further validated using sequencing from the same cell, as opposed to scRNA-seq from one cell being compared to exome sequencing from a bulk of cells as was performed with the articular chondrocytes. This would also address the shortcomings tied with validation via Sanger sequencing which uses material from a bulk of cells.

## 6. Conclusion

Based on the experimental and simulated data, Red Panda provides a distinct advantage over other available software. This improvement comes from its ability to more accurately predict homozygous-looking and bimodally-distributed heterozygous variants as compared to other tools. Due to the unique nature of scRNA-seq data, one must treat heterozygous variants with special consideration, and Red Panda provides a custom approach to this class of variants. From these results, it is clear that due to the inherent nature of RNA expression patterns in single cells, it is difficult to assess what variants exist with the same accuracy that we can with standard exome or WGS. Despite this, Red Panda provides a novel method of identifying variants in scRNA-seq and performs this function better than variant callers designed for bulk NGS datasets.

## Authors’ contributions

AC designed the study, performed all analyses, and wrote the manuscript. SR, KS, and SP performed experiments. CG with the help of NKM supervised the project, provided essential feedback on experimental results and input on the analyses themselves, and thoroughly edited the manuscript. KB and AD provided technical guidance and materialistic support for experiments performed by SR/SP and KS, respectively. All authors read and approved the final manuscript.

## Acknowledgements

The authors are very grateful to Kristin Wipfler for her valuable input during algorithm development, Cecily Zdan for copy editing, and Dr. James Eudy for his valuable input during setup and processing of the sequencing.

## Funding

This work was supported by the development funds to CG, and the Office of Graduate Studies fellowship to AC, from the University of Nebraska Medical Center (UNMC) and the NIH subaward [2P01AG029531] to CG. The University of Nebraska DNA Sequencing Core and the Bioinformatics and Systems Biology core receive partial support from the National Institutes of Health grants [P20GM103427, 1P30GM110768, P30CA036727].

## Conflict of Interest

The authors declare that the research was conducted in the absence of any commercial or financial relationships that could be construed as a potential conflict of interest.

## Notes

#### Summary of Updates

Changing the publication category from 'Confirmatory Results' to 'New Results'

