## Supplementary data for "Red Panda: A novel method for detecting variants in single-cell RNA sequencing"

**Supplemental figures and data**

**Figures**

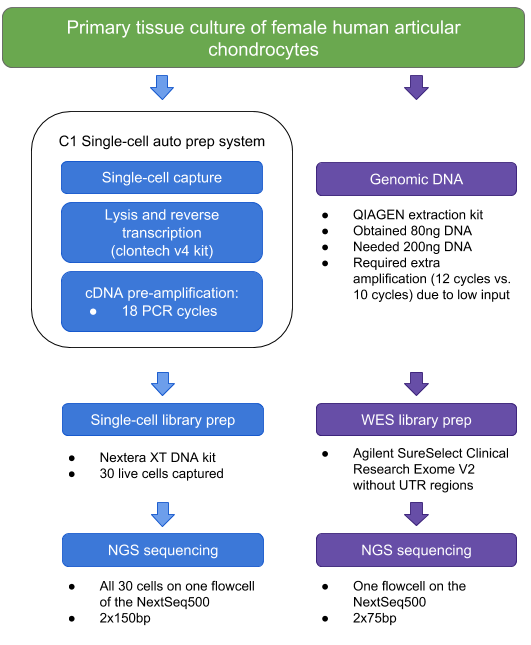

**Supplemental Figure 1. Human articular chondrocyte sequencing strategy.** Exome sequencing was paired with scRNA-seq for the the primary tissue culture of human articular chondrocytes. The library prep for the single cells was performed using the updated Smart-seq2 protocol.

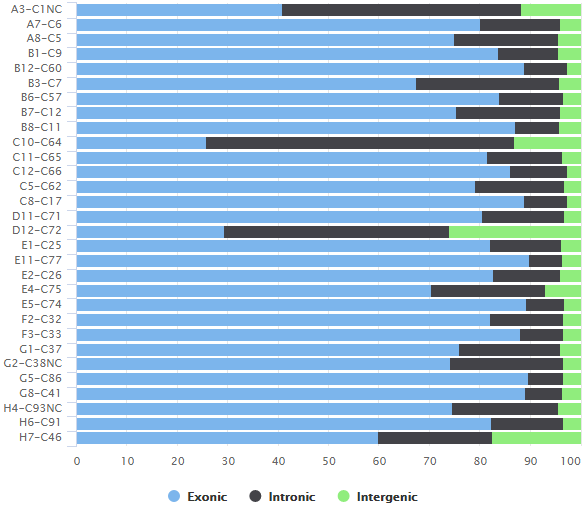

**Supplemental Figure 2. The genomic origin of reads found in each cell.** Here one can see what percentage of reads originate from exons (blue), introns (black) or intergenic space (green). The cells A3-C1NC, C10-C64, D12-C72, and H7-C46 have significantly more reads originating outside the exonic region than other samples.

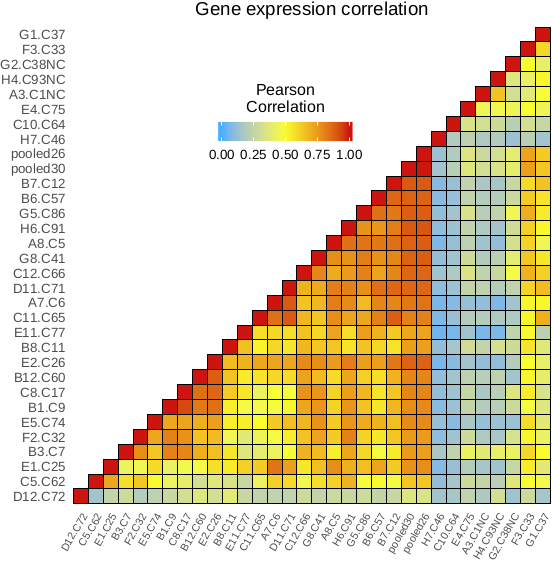

**Supplemental Figure 3. Expression correlation between articular chondrocytes.** Pearson Correlation Coefficient calculated for every possible comparison of cells to each other and the two batches of cells. The darker the color red, the higher the correlation between each cell. “pool26” contains reads pooled from 26 cells, A3-C1NC, C10-C64, D12-C72, and H7-C46 were removed; “pool30” contains reads pooled from all 30 cells. as well as identify three other cells that do not correlate well based on their expression patterns: H4-C93NC, G2-C38NC, and E4-C75.

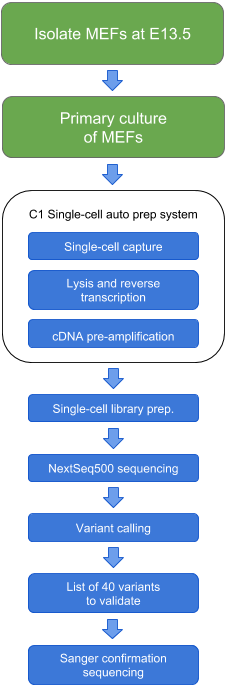

**Supplemental Figure 4. The sequencing strategy for the MEFs.** The MEFs have variant calling performed on them with five variant callers as with the articular chondrocytes. Validation is performed by Sanger sequencing on 40 variants as well as using simulations based on this sequencing data.

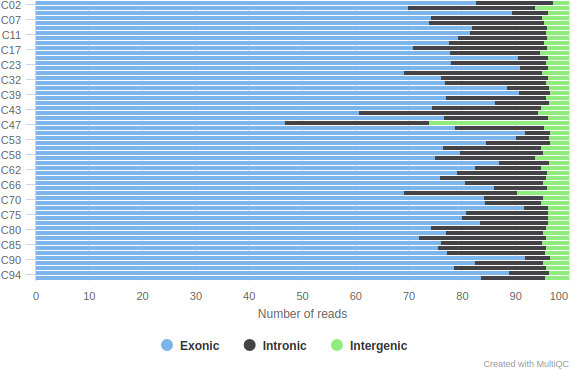

**Supplemental Figure 5. The genomic origin of reads found in each MEF.** Here one can see what percentage of reads originate from exons (blue), introns (black) or intergenic space (green). The cell C47 is the only cell to have significantly more reads originating outside the exonic region than other samples**.**

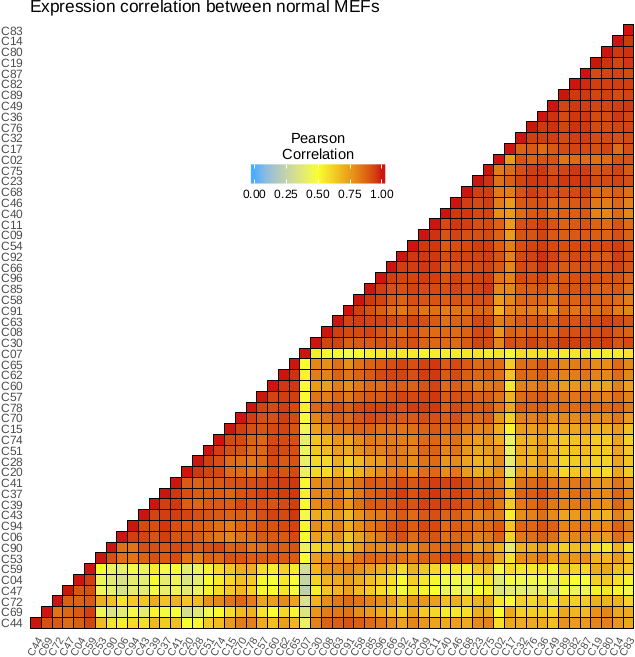

**Supplemental Figure 6. Expression correlation between normal MEFs.** Pearson Correlation Coefficient calculated for every possible comparison of cells to each other for the normal MEFs. The darker the color red, the higher the correlation between each cell. One can clearly see one cell that fails to correlate will with the other cells: C07. The bottom block of cells significantly correlates with a high number of cells and they are therefore retained.

| **a** | 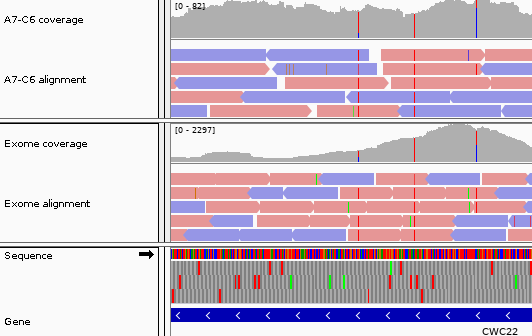 |
| --- | --- |
| **b** | 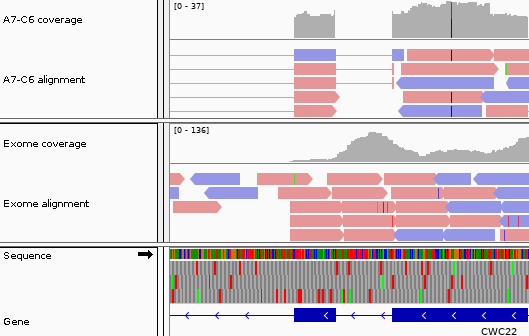 |
| **Supplemental Figure 7.** Proof of concept data in articular chondrocytes. An example of the variations, from gene CWC22, that we find in the scRNA-seq data as compared to the exome. The main area of interest is the coverage track (the gray histograms). Red corresponds to T and blue corresponds to a C. When there are two colors, the top color corresponds to the alternate allele. **(a)** Two hetSNVs found in the cell A7-C6 have reads supporting them at percentages of 80% (left) and 20% (right). The same hetSNVs are found in the exome data at 50%. There is also a homozygous variant (middle) seen in both. **(b)** One hetSNV found in the same gene at 53% in the cell A7-C6 is absent in the exome sequencing. This is expected as it does not fit the existing biomodal distribution at 80% or 20%.  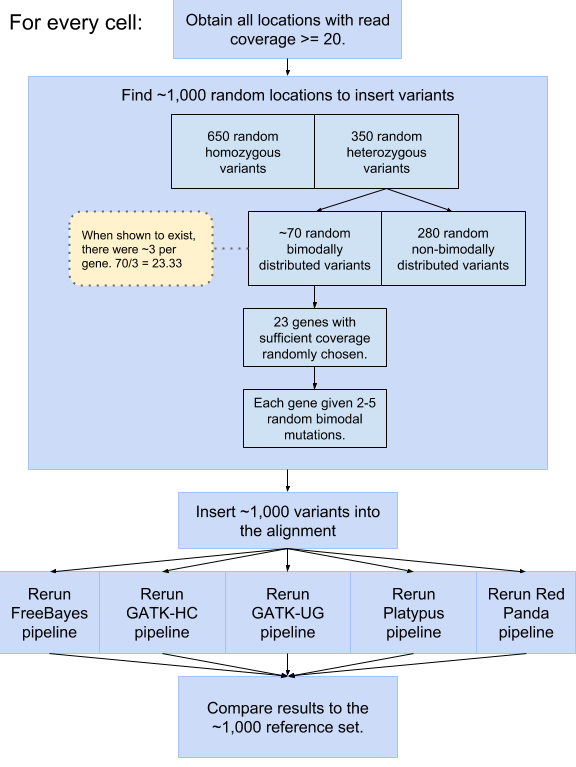  **Supplemental Figure 8**. **Workflow for inserting simulated variants.** To assess each tool, ~1,000 simulated variants (650 homozygous, 280 heterozygous, and ~70 bimodally-distributed heterozygous) were inserted into the alignments for each cell. Standard variant calling was then performed using each tool, and these results were compared to the list of known variants to assess their performance. | |

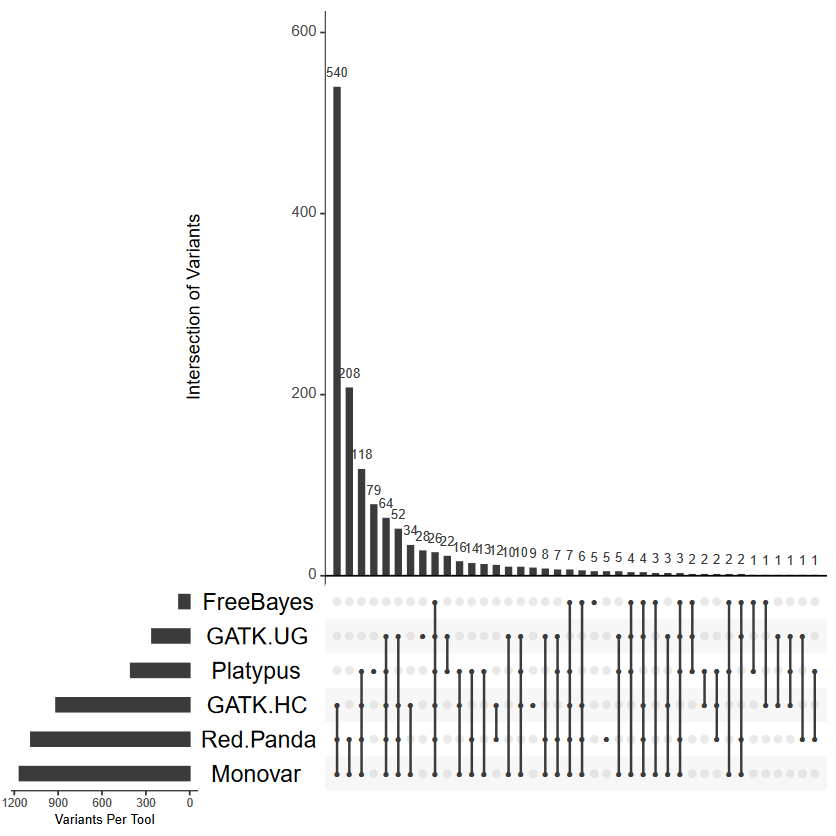

**Supplemental Figure 9. UpSet plots of the overlap between each tool.** The overlap of the variants identified by each tool can be seen for the cell G1-C37. Each column of the X-axis shows the overlap between each tool represented by a filled-in dot. For example, the first column indicates that GATK-HC, Monovar, and Red Panda shared 540 variants, the second shows that Red Panda and Monovar share 208 variants, the third column indicates that there were 118 variants shared between Platypus, GATK-HC, Monovar, and Red Panda, and so on.

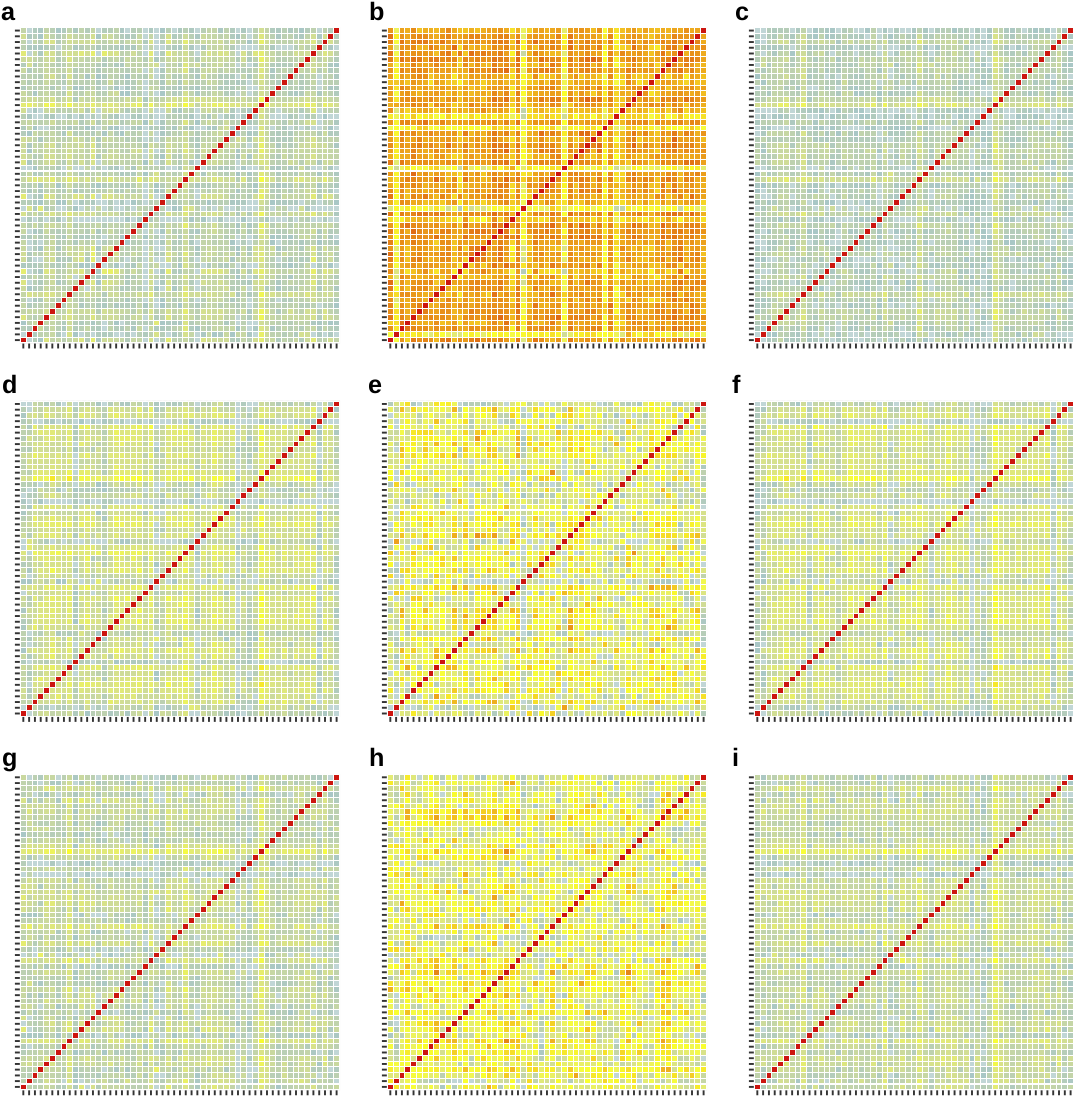

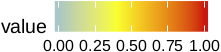

**Supplemental Figure 10. The fraction of overlap in variants for every cell using FreeBayes, GATK HC, and GATK UG.** The fraction of overlap for **(a-c)** FreeBayes, **(d-f)** GATK-HaplotypeCaller, and **(g-i)** GATK-UnifiedGenotyper when comparing **(a, d, g)** all variants, **(b, e, h)** homozygous-looking variants, and **(c, f, i)** heterozygous variants. Each box in the matrix is a comparison between two cells.

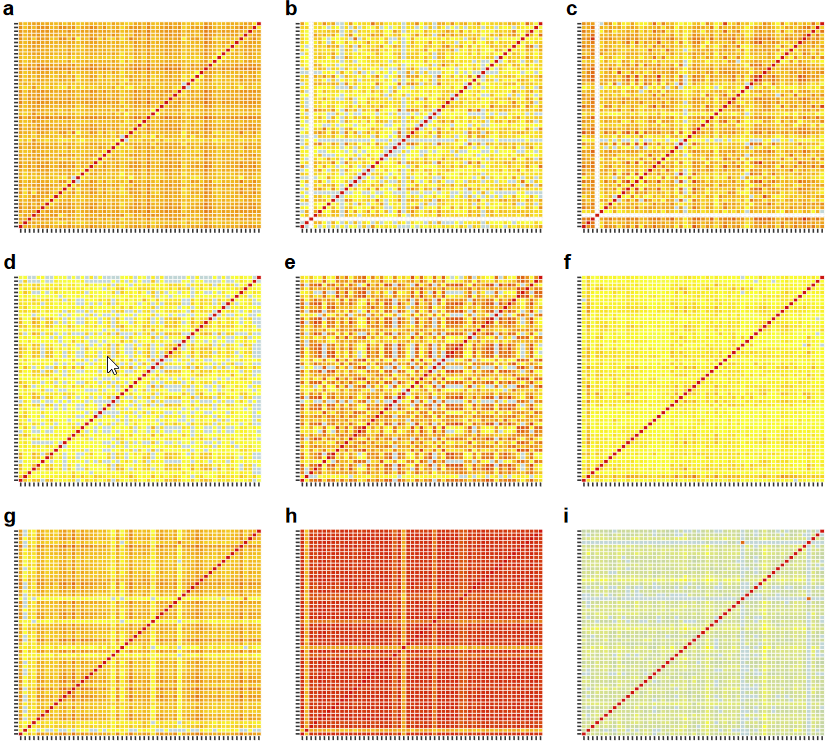

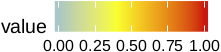

**Supplemental Figure 11. The fraction of overlap in variants for every cell using Monovar, Platypus, and Red Panda.** The fraction of overlap for **(a-c)** Monovar, **(d-f)** Platypus, and **(g-i)** Red Panda when comparing **(a, d, g)** all variants, **(b, e, h)** homozygous-looking variants, and **(c, f, i)** heterozygous variants. Each box in the matrix is a comparison between two cells.

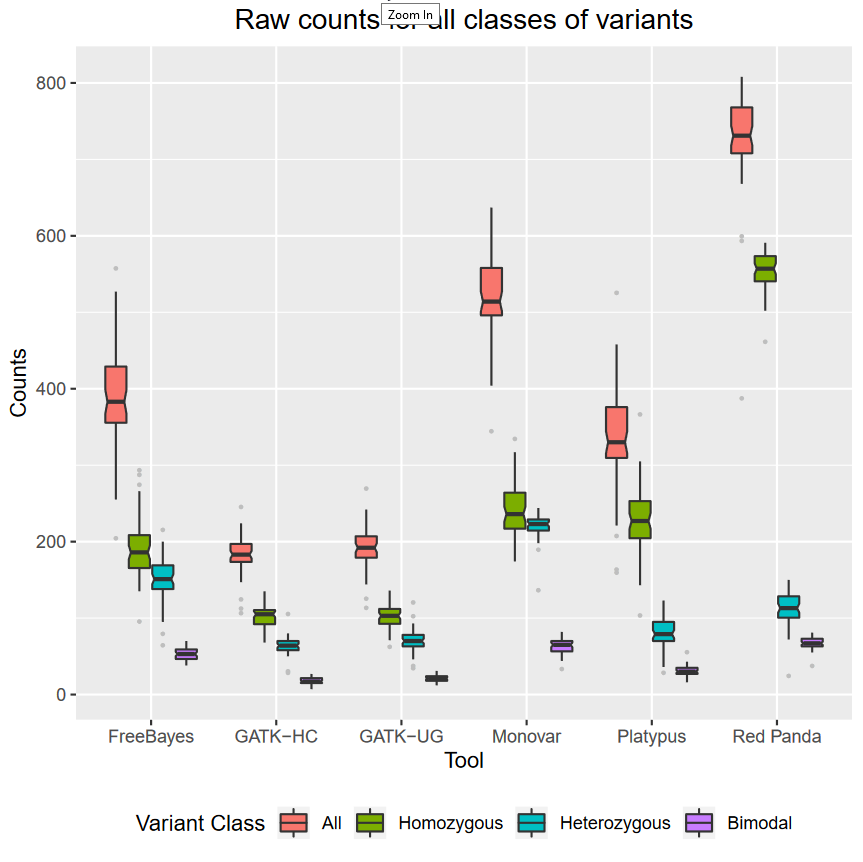

**Supplemental Figure 12. Raw counts of True Positives for each tool.** The box plots of the raw number of True Positives show how well each tool is at identifying variants in the simulation. Due to advantages gained in identifying homozygous and bimodally-distributed variants, Red Panda identifies the highest number of True Positives.


### Tables

**Supplemental Table 1. Eight human articular chondrocytes removed for quality reasons. “Too many reads outside exon” is defined as one standard deviation above the median percentage of reads aligned outside exons for all samples. Statistically insignificant correlation coefficient is defined as p > 0.05 for a Pearson correlation coefficient when comparing the transcription profile of a single cell to the pool of all 30 cells.**

| **Sample Name** | **Reason for removal** |
| --- | --- |
| A3-C1NC | Too many reads outside exon; Statistically insignificant correlation coefficient |
| C10-C64 | Too many reads outside exon; Statistically insignificant correlation coefficient |
| C12-C66 | Low read count: 80k total reads. |
| D12-C72 | Too many reads outside exon; Statistically insignificant correlation coefficient |
| E4-C75 | Statistically insignificant correlation coefficient |
| G2-C38NC | Statistically insignificant correlation coefficient |
| H4-C93NC | Statistically insignificant correlation coefficient |
| H7-C46 | Too many reads outside exon; Statistically insignificant correlation coefficient |

**Supplemental Table 2. Parameters used to design the primers used for PCR and Sanger.** Parameters with a * were changed from their defaults to ensure good sequencing.

| **Parameters used to design primers on Primer3Plus** | |
| --- | --- |
| Product Size Range | 401-700* |
| Min primer | 18 |
| Opt primer* | 20 |
| Max primer | 27 |
| Primer Tm Min | 57 |
| Primer Tm Opt | 60* |
| Primer Tm Max | 63 |
| Max Tm difference | 100 |
| Primer GC% Min | 20 |
| Primer GC% Opt | 50* |
| Primer GC% Max | 80 |
| Concentration of monovalent cations | 50 |
| Concentration of divalent cations | 0 |
| Annealing Oligo Concentration | 50 |
| Concentration of dNTPs | 0 |
| Max Self Complementarity | 4* |
| Max #Ns | 0 |
| Max Poly-X | 5 |
| CG Clamp | 1* |
| Max 3' Self Complementarity | 3 |
| Max 3' Stability | 9 |
| Pair Max Repeat Mispriming | 24 |
| Pair Max Template Mispriming | 24 |

**Supplemental Table 3. Human articular chondrocyte exome sequencing statistics.** Sequencing and analysis statistics of the exome data from the human articular chondrocytes.

| **Total Reads** | **Paired Reads** | **PCR Duplicate** | **Alignment** | **Coverage** | **On-target rate** |
| --- | --- | --- | --- | --- | --- |
| 182M | 111M (61%) | 38.30% | 98.70% | 74.67x | 55% |

**Supplemental Table 4. Human articular chondrocyte exome variant calling statistics.** Variant analysis statistics of the exome data from the human articular chondrocytes using the ensemble approach where 2/3 variant caller tools had to agree to call a variant.

| **Bed file contains 100bp ± coding exons** | **Total Variants** | **Homozygous Variants** | **Heterozygous Variants** | **Ratio** | **SNVs** | **indels** |
| --- | --- | --- | --- | --- | --- | --- |
| Yes | 85,128 | 33,856 | 45,769 | 0.74 | 79,627 | 5,504 |
| No | 20,315 | 7,777 | 12,538 | 0.62 | 20,057 | 258 |

**Supplemental Table 5.** **Summary of the human articular chondrocytes captured on the C1.**

| **Live Cells** | **Dead Cells** | **No Color (NC)** | **Live and Dead** | **>1 Live** | **>1 Dead** | **Empty** |
| --- | --- | --- | --- | --- | --- | --- |
| 27 | 37 | 3 | 17 | 2 | 3 | 7 |

**Supplemental Table 6.** **Summary table of variants identified by Red Panda in human articular chondrocytes**. Percent of variants that are homozygous-looking, heterozygous, heterozygous and validated by GATK-HC, heterozygous and validated by Red Panda are calculated. Average total number of variants in the final VCF file is also shown.

|  | **Total** |
| --- | --- |
| **Percent of total that are homozygous** | 69.78% |
| **Percent of total that are heterozygous** | 30.22% |
| **Percent of Heterozygous from RP** | 77.17% |
| **Percent of Heterozygous from GATK-HC** | 22.83% |
| **Total Variants** | 1369.5 |

**Supplemental Table 7. Validation of variants identified by all five variant callers.** In the Validated by Sanger column, N = No, and NS = No Sequence at that position.

| **Number** | **Hom/Het** | **Variant** | **Variant** | **Validated by Sanger** | **Cells supported by** |
| --- | --- | --- | --- | --- | --- |
| 1 | Het | chr1 43954701 | T→G | N | 2 |
| 2 | Hom | chr1 181176175 | C→T | N | 2 |
| 3 | Het | chr2 3328501 | G→T | N | 2 |
| 4 | Het | chr2 22940605 | G→C | NS | 2 |
| 5 | Het | chr2 33246775 | A→G | N | 2 |
| 6 | Het | chr2 39195366 | A→G | N | 2 |
| 7 | Het | chr3 19133919 | A→G | N | 2 |
| 8 | Het | chr4 43977653 | A→G | N | 2 |
| 9 | Het | chr8 36567823 | T→C | N | 2 |
| 10 | Het | chr8 71359979 | A→G | N | 2 |
| 11 | Het | chr6 83802489 | G→T | N | 2 |
| 12 | Het | chr9 44742670 | C→A | N | 2 |
| 13 | Het | chr10 112926193 | T→C | N | 2 |
| 14 | Hom | chr11 73175960 | A→G | N | 2 |
| 15 | Het | chr13 75771943 | G→T | NS | 2 |
| 16 | Het | chr13 90105223 | T→C | N | 2 |
| 17 | Het | chr16 49868008 | C→T | N | 2 |
| 18 | Hom | chr16 58466497 | G→A | N | 2 |
| 19 | Het | chr17 12683939 | A→G | N | 2 |
| 20 | Het | chr18 43321798 | T→C | N | 2 |

**Supplemental Table 8. First sequencing pass: Validation of variants only identified by Red Panda.** In the Validated by Sanger column, Y = Yes, N = No, and NS = No Sequence at that position.

| **Number** | **Hom/Het** | **Variant Location** | **Variant** | **Validated by Sanger** | **Cells supported by** |
| --- | --- | --- | --- | --- | --- |
| 1 | Het | chr2 120515974 | T→C | N | 2 |
| 2 | Het | chr3 19133919 | A→G | N | 2 |
| 3 | Hom | chr3 95734876 | T→C | NS | 38 |
| 4 | Het | chr4 130165817 | T→C | N | 2 |
| 5 | Hom | chr4 132833055 | C→G | NS | 9 |
| 6 | Hom | chr5 104435120 | C→G | NS | 2 |
| 7 | Hom | chr7 27205154 | TA→T | NS | 2 |
| 8 | Hom | chr7 27205568 | A→G | NS | 4 |
| 9 | Het | chr8 85261271 | A→C | NS | 2 |
| 10 | Het | chr8 85261288 | G→A | NS | 2 |
| 11 | Hom | chr10 40251185 | G→A | N | 2 |
| 12 | Hom | chr11 72777865 | C→A | NS | 2 |
| 13 | Hom | chr12 54783425 | T→C | NS | 2 |
| 14 | Het | chr13 31630905 | A→G | N | 2 |
| 15 | Het | chr14 54542219 | T→C | NS | 2 |
| 16 | Het | chr16 52270742 | C→A | N | 2 |
| 17 | Hom | chr16 94468834 | C→T | Y | 34 |
| 18 | Het | chr19 60771023 | C→A | NS | 2 |
| 19 | Het | chr19 60771042 | G→T | NS | 2 |
| 20 | Hom | chrX 101404519 | C→A | N | 2 |

**Supplemental Table 9. Second sequencing pass: Validation of variants only identified by Red Panda.** In the Validated by Sanger column, Y = Yes, N = No, and NS = No Sequence at that position.

| **Number** | **Hom/Het** | **Variant Location** | **Variant** | **Validated by Sanger** | **Cells supported by** |
| --- | --- | --- | --- | --- | --- |
| 1 | Het | chr2 120515974 | T→C | N | 2 |
| 2 | Het | chr3 19133919 | A→G | N | 2 |
| 3 | Hom | chr3 95734876 | T→C | Y | 38 |
| 4 | Het | chr4 130165817 | T→C | N | 2 |
| 5 | Hom | chr4 132833055 | C→G | Y | 9 |
| 6 | Hom | chr5 104435120 | C→G | N | 2 |
| 7 | Hom | chr7 27205154 | TA→T | NS | 2 |
| 8 | Hom | chr7 27205568 | A→G | N | 4 |
| 9 | Het | chr8 85261271 | A→C | N | 2 |
| 10 | Het | chr8 85261288 | G→A | NS | 2 |
| 11 | Hom | chr10 40251185 | G→A | N | 2 |
| 12 | Hom | chr11 72777865 | C→A | NS | 2 |
| 13 | Hom | chr12 54783425 | T→C | NS | 2 |
| 14 | Het | chr13 31630905 | A→G | N | 2 |
| 15 | Het | chr14 54542219 | T→C | NS | 2 |
| 16 | Het | chr16 52270742 | C→A | N | 2 |
| 17 | Hom | chr16 94468834 | C→T | Y | 34 |
| 18 | Het | chr19 60771023 | C→A | N | 2 |
| 19 | Het | chr19 60771042 | G→T | N | 2 |
| 20 | Hom | chrX 101404519 | C→A | N | 2 |
